# Boosting polygenic risk scores

**DOI:** 10.1101/2022.04.29.489836

**Authors:** Hannah Klinkhammer, Christian Staerk, Carlo Maj, Peter M. Krawitz, Andreas Mayr

## Abstract

Polygenic risk scores (PRS) evaluate the individual genetic liability to a certain trait and are expected to play an increasingly important role in the field of clinical risk stratification. Most often, PRS are estimated based on summary statistics of univariate effects derived from genome-wide association studies. To improve the predictive performance of PRS, it is desirable to fit multivariable models directly on the genetic data. Due to the large and high-dimensional data, a direct application of existing methods is often not feasible and new efficient algorithms are required to overcome the computational burden regarding efficiency and memory demands.

We develop an adapted component-wise *L*_2_-boosting algorithm to fit genotype data from large cohort studies to continuous outcomes using linear base-learners for the genetic variants. Similar to the snpnet approach implementing lasso regression, the proposed snpboost approach iteratively works on smaller batches of variants. By restricting the set of possible base-learners in each boosting step to variants most correlated with the residuals from previous iterations, the computational efficiency can be substantially increased without losing prediction accuracy. Furthermore, for large-scale data based on various traits from the UK Biobank we show that our method yields competitive prediction accuracy and computational efficiency compared to the snpnet approach. Due to the modular structure of boosting, our framework can be further extended to construct PRS for different outcome data and effect types.

## Introduction

In times of next generation sequencing and decreasing costs for whole genome sequencing, the amount of available genotype data has increased dramatically in recent years, giving rise to new genetic insights [1, 2].

Polygenic risk scores (PRS) measure the individual genetic liability to a certain trait and can provide relevant information in the context of disease-risk stratification. In contrast to high-impact monogenic variants, which are mostly rare and have a high effect size, PRS are derived from common variants such as single-nucleotide polymorphisms (SNPs) with low or medium effect sizes. Polygenic effects could also explain part of the incomplete penetrance seen in many identified monogenic variants, as for example in the genes BRCA1 and BRCA2 both leading to a highly increased risk of breast cancer [3]. Recent studies on the UK Biobank suggest that high-impact monogenic variants, PRS and family history could contribute additively to the risk of developing breast and prostate cancer [4]. Despite these findings, PRS still lack to explain relevant parts of the estimated heritability of many traits.

PRS are typically derived as a weighted sum of univariate effect estimates of the measured variants based on summary statistics from genome-wide association studies (GWAS) [5]. Despite several approaches to account for linkage disequilibrium (LD, referring to the correlation structure between variants) and for the selection of informative variants [6–9], the univariate structure of the estimation cannot fully account for interdependencies between the variants. A natural extension of using univariate models could be to fit a single multivariable model. While this approach seems natural from a methodological perspective, a direct application of existing methods is typically infeasible due to the high dimensionality of the genotype data, which can easily exceed the available computer memory. Recently, some approaches have been proposed to overcome this computational burden [10–12]. In particular, Qian et. al. proposed the so-called batch screening iterative lasso (BASIL) algorithm to fit the lasso on the complete original genotype data [11, 13]. The algorithm works on subsets of variants and computes the complete lasso path in an iterative fashion. Apart from the lasso, the algorithm can also be extended to other penalized regression methods such as the relaxed lasso [14] or the elastic net [15]. In this context, Qian et al. were able to demonstrate that multivariable regularized PRS models fitted via the BASIL algorithm outperform the classical GWAS-based PRS for various traits such as height and high cholesterol.

While penalized regression models like the lasso and the elastic net impose explicit regularization, statistical boosting represents an alternative approach by introducing an implicit algorithmic regularization when combined with early stopping of the algorithm [16, 17]. Boosting algorithms iteratively fit pre-defined base-learners to the gradient of the loss function, selecting the most influential base-learner in each step. The main tuning parameter of boosting algorithms is the number of iterations, which enables implicit variable selection and leads to sparse models. Due to its modular structure, boosting allows to combine possible base-learners with any convex loss function. These algorithms hence offer a great flexibility in the field of statistical modelling, including various response types and the estimation of non-linear or other types of effects. A recent work has incorporated boosting into PRS modelling via a three-step approach [12]: First, a marginal screening approach was applied on all variants to identify potentially informative ones. Then, multivariable algorithms including probing with boosting [18] were applied on blocks of variants in LD to select (“fine-map”) the most informative variants. Finally, a statistical boosting model was fitted on the variant set created by joining the selected variants of all chunks. While this approach yielded particularly sparse and interpretable models, their predictive performance was comparable to PRS derived by univariate methods like clumping and thresholding [6] and was outperformed by the predictive performance achieved by the lasso via the BASIL algorithm.

In this article we introduce a new framework to boost PRS, starting with a new computational approach to build *L*_2_-boosting models on large-scale genotype data for quantitative traits. Similar to the snpnet approach for the lasso, our algorithm iteratively works on smaller batches of variants. Yet, in contrast to recent boosting methods [12, 19], the variants do not need to be pre-filtered in our snpboost approach and the batches are not pre-defined or randomly sampled, but chosen iteratively and deterministically in a data-driven way based on the correlations of the variants to the remaining residuals. By restricting the set of available base-learners in each step to those variants which were most correlated with residuals from a previous iteration, we are able to reduce the search space and decrease the computational time compared to a classical component-wise boosting algorithm. We conducted a simulation study to examine the performance of our adapted boosting algorithm snpboost compared to the original *L*_2_-boosting on a reduced but still high-dimensional data set, on which the application of standard *L*_2_-boosting was still computationally feasible. Furthermore, we simulated data of higher dimensionality and larger sample size to investigate the influence of various hyperparameters (including the batch size) on the prediction accuracy and computational burden of the snpboost approach in a typical large-scale setting. We discuss reasonable default values for the hyperparameters which are incorporated in the provided R implementation (https://github.com/hklinkhammer/snpboost). Finally, we constructed multivariable PRS for various traits on data from the UK Biobank via application of snpboost and compared the performance of our approach to the lasso estimates from the BASIL algorithm proposed by Qian et al. On the examined phenotypes we found highly comparable predictive performance while our adapted boosting approach had a tendency to select sparser models compared to the lasso.

## Methods

For 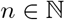 individuals, let 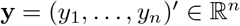 denote a particular continuous phenotype of interest. Furthermore, let *X_j_* correspond to the genetic variant *j*, for *j* = 1,…, *p*. The observed dosage data of *n* individuals is given in the genotype matrix **X** = (*x_i,j_*) ∈ [0, 2]*^nxp^*, where **x***_j_* ∈ [0, 2]*^n^* corresponds to the *j*-th column of **X**. We consider a linear regression model

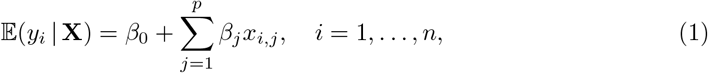

with coefficients 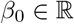 and 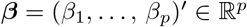. The aim is to determine coefficients 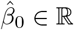 and 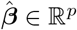 such that the estimator

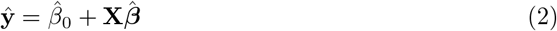

minimizes the mean squared error of prediction on an independent test set

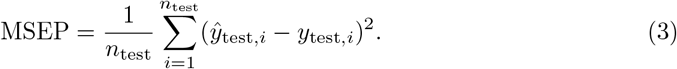

Additionally, one is often interested in relatively sparse models in the sense that only a fraction of the coefficient vector 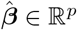 is non-zero.

In high-dimensional settings with *p* > *n* it is not feasible to apply classical estimation techniques like the ordinary least squares estimator. A commonly-used solution is to consider further constraints on the coefficient vector resulting in penalized regression methods including the lasso [13]. The lasso incorporates an *L*_1_-penalty on the coefficient vector such that the lasso estimate 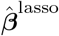 is given by

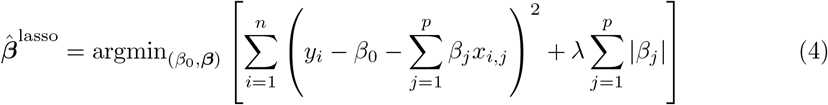

for some *λ* ≥ 0. The explicit *L*_1_-penalization of the coefficient vector leads to shrinkage of the coefficient estimates. In contrast to ridge regression [21], the use of the *L*_1_-penalty enables to set some parameters exactly to zero corresponding to sparse models. There has been extensive research on the theoretical properties of the lasso including oracle inequalities in high-dimensional settings (e.g. [22–25]). Nevertheless, there are situations leading to variable selection problems of the lasso, particularly in the presence of high correlations between signal and noise variables [26]. When working with genotype data, high correlations between signal and noise variables might often be present as a result of linkage desequilibrium (LD), i.e. genetic variants that have close positions on the DNA strand tend to be highly correlated.

An alternative to explicitly penalized regression methods such as the lasso is statistical gradient boosting [16, 17]. Gradient boosting requires the specification of a loss function 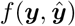 and so-called base-learners *h_j_* that are iteratively fitted to the response. In detail, the aim is again to fit the linear regression model (1) which is performed in an iterative fashion. Starting at iteration *m* = 0 with a starting value 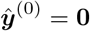, the following steps are repeated until a maximum number *m*_stop_ of boosting iterations is reached [16]:

1. Set *m*: = *m* +1 and compute the negative gradient vector of the loss function:

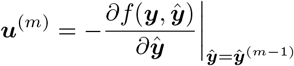
2. Fit every base-learner *h_j_* separately to the negative gradient vector ***u***^(*m*)^ and select the best fitting base-learner 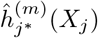.
3. Update the predictor with the learning rate 0 ≤ *ν* ≤ 1:

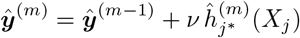
4. Stop if *m* = *m*_stop_.

Stopping the algorithm before it converges (early stopping) leads to implicit regularization and shrinkage of effect estimates. The component-wise *L*_2_-boosting algorithm [16, 27] employs the squared error 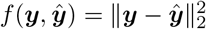 as a loss function [27] and separate univariate linear regression models of the outcome on the j-th genetic variant as base-learners (i.e. *h_j_*(*X_j_*) = *β*_0_ + *β_j_X_j_*, for *j* = 1,…, *p*). In low-dimensional (*p* < *n*) settings this set-up mimics a classical Gaussian linear model and converges to the least squares solution for large values of *m*_stop_. The general boosting procedure can be interpreted as gradient descent in function space, where the residual vector represents the gradient of the *L*_2_ loss and the function space is provided by the different base-learner solutions [17, 27]. The previously described steps transform therefore into the following procedure (shown in grey in Figure 1):

**Fig 1.**
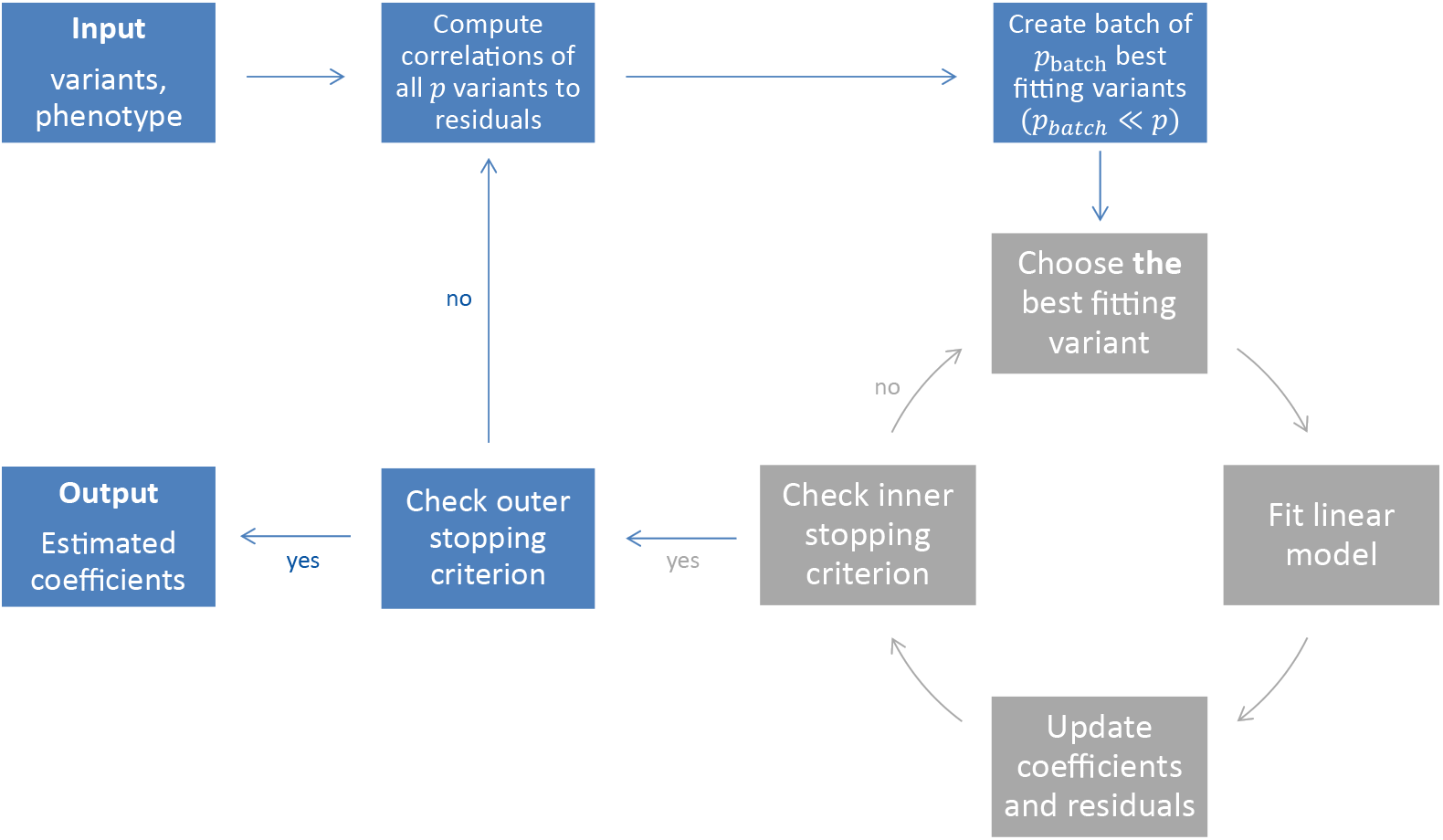
Illustration of the snpboost algorithm. The snpboost algorithm consists of an outer loop to create batches (shown in blue) and an inner loop representing the boosting on one batch (shown in grey).

The best fitting base-learner in boosting step *m* + 1 corresponds to the variant *j*^*^ with the highest Pearson correlation *ρ*(***x**_j*_*, ***r***^(*m*)^) to the residuals 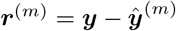 resulting from the previous boosting step *m*. We then fit a linear regression model of the current residuals ***r***^(*m*)^ on the variant *j*^*^ and update the corresponding coefficient 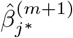 as well as the intercept 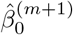. This is repeated until a maximum number of boosting iterations is reached or any other early stopping criterion is fulfilled. If additional covariates apart from the genetic variants are included in the model, they are treated as mandatory covariates – similar to the intercept. The additional covariates are included in each single base-learner and are hence updated in each boosting step without competing with the genetic variants.

Hepp et al. [26] investigated the commonalities and differences between the lasso and statistical boosting: while there are (low-dimensional) settings in which the gradient boosting approximates the lasso coefficient paths arbitrarily close when the learning rate *ν* is approaching 0, their results generally differ if the coefficient paths are not monotone. The authors note that, in contrast to the lasso which limits the sum of the absolute values of the coefficients for each penalty parameter *λ* separately, boosting limits the total *L*_1_-arc-length of all coefficient curves [26]. Interpreting this as the total absolute distance “travelled” by all coefficients among the coefficient paths through the iterations *m* = 1,…, *m*_stop_, it becomes clear that the solution in a certain iteration depends on all previous solutions of the iterative algorithm. This might lead to more stable pathways particularly in settings with high correlations between independent variables, which is typical for genetic data. Hepp et al. conducted several numerical experiments including high-dimensional settings in which they found similar predictive performance of lasso and boosting. In detail, boosting tended to yield slightly better prediction results while the lasso tended to result in sparser models with faster computations. On the other hand, the boosting algorithm can be easily extended to different response types as well as to different effects, including non-linear and interaction effects.

When working on genetic data from large cohort studies we do not only face a high-dimensional setting with *p* > *n* but also a large-scale setting with large sample sizes *n and* large numbers of variants *p*. Large-scale settings often lead to extended computational times as well as memory issues. To overcome these and apply statistical boosting on genotype data, we implemented an adapted component-wise *L*_2_-boosting algorithm that is built on the snpnet framework [11] and works on batches of variants. To do so, we additionally incorporate a batch-building step before starting the boosting iterations (shown in blue in Fig 1). In this step we extract the *p*_batch_ variants (*p*_batch_ ≪ *p*) with the highest correlation *ρ*(***x**_j_*, ***r***^(*m*)^) to the current residual vector and include them in the batch *B_k_*. A maximum number of *m*_batch_ boosting iterations is performed on batch *B_k_* before the next batch is built based on the correlations of all *p* variants to the updated residuals. In total, we fit a maximum of *b*_max_ batches or stop early if an early stopping criterion is fulfilled. The algorithm is summarised in Table 1 and Fig 1.

**Table 1.**
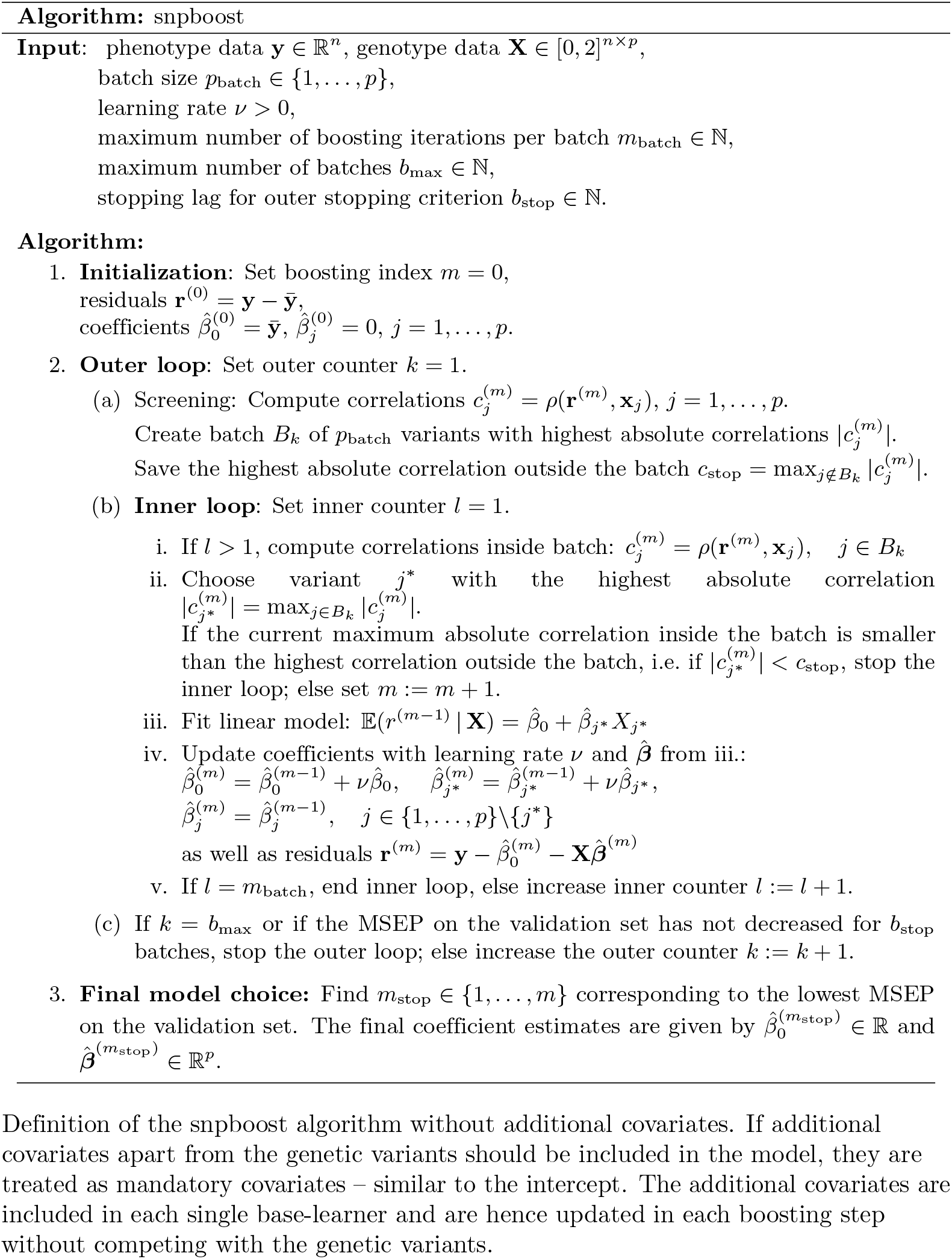
Definition of the snpboost algorithm.

By iteratively working on batches of variants we save computational time and memory because only parts of the variants have to be loaded into memory at once. Additionally, not every step requires the calculation of all potential base-learner solutions and the updated correlations for all variants. By this, we encourage additional sparsity by restricting the search space in terms of the set of available base-learners (as variants not included in the current batch cannot be selected). To examine when a new set of base-learners should be considered, which corresponds to the question when to stop the inner loop (inside the batches) and create a new batch of variants, we incorporated another step: we monitor the correlations of the variants inside a batch to the residuals and compare them to the correlations of variants outside of the batch. When creating a batch *B_k_* we therefore compute and store the highest outer correlation

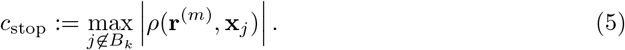

After each boosting step *m* we check if the greatest absolute correlation of the variants inside the batch *B_k_* to the current residual vector **r**^(*m*)^ is smaller than *c*_stop_:

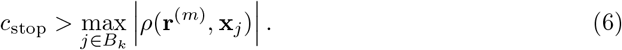

If inequality (6) holds true, we stop the inner loop and create a new batch since a variant outside the batch may provide a better fit to the current residual vector. In the original *L*_2_-boosting without batches, the variant with the highest correlation to the residuals would be chosen in each boosting step. The incorporation of batches in general limits this choice to the variants inside the batch. However, the proposed stopping criterion provides an indication to consider variants outside the batch which may be higher correlated with the current residuals. Actually, if all variants were independent, the proposed stopping criterion would lead to the same choice of variants in each boosting step in snpboost as in the original *L*_2_-boosting. Despite linkage disequilibrium, our simulation results show that the proposed stopping criterion yields reasonable variant choices and results in a competitive predictive performance (Section Comparison to original *L*_2_-boosting in smaller settings). Additionally, the inner loop is also stopped if a pre-specified maximum number of *m*_batch_ inner boosting iterations have been performed on the same batch (i.e. the number of updates inside the batch reaches *m*_batch_).

Furthermore, we need to determine after how many batches the algorithm should terminate. In classical statistical boosting the number of boosting iterations is often selected by cross-validation or resampling techniques – mimicking an additional data set to validate the predictive performance of the resulting models. However, if the data set is large enough, one can also directly divide the data into training and validation set. As in Qian et al. [11], we hence simultaneously monitor the predictive performance of our model on an independent validation set while fitting on the training set. As a validation criterion for the predictive performance we use the mean squared error of prediction on the validation set. The outer loop consisting of the batch-building step is stopped if the MSEP on the validation set has not decreased for *b*_stop_ batches or after a maximum number of *b*_max_ batches have been processed.

The proposed method is implemented as an add-on to the `snpnet` package by Qian et al. [11] in the statistical computing environment R ([28], https://github.com/hklinkhammer/snpboost). While we are also incorporating PLINK 2.0 [29] to compute the correlations and build the batches in the outer loop, we replaced the fitting of the lasso by the adapted component-wise *L*_2_-boosting algorithm on the resulting batches (see Table 1 and Fig 1).

## Simulation Study

We conducted a simulation study to investigate the behaviour of the proposed snpboost algorithm in various controlled data scenarios. The simulation study aims at two main goals: first, to examine potential differences in performance compared to the original component-wise *L*_2_-boosting [27] in smaller settings and, second, to gain insights on how to choose the included hyperparameters in practical situations.

Simulations are based on the UK Biobank genotype data [30] obtained under application number 81202 combined with simulated phenotypes. We restricted the individuals to white British ancestry and used the PLINK 2.0 function --`thin-indiv-count` to randomly sample *n* individuals, of which 50%, 20% and 30% were assigned to the training, validation and test set, respectively ([29], [31]). Then, *p* variants with minor allele frequency not less than 1 % were randomly sampled using PLINK 2.0’s --`thin-count`. Missing genotypes were replaced by the reference allele using the R package `bigsnpr` [10].

Continuous phenotypes were simulated from a linear model with Gaussian distributed noise and effect sizes using `bigsnpr`. To account for different genetic architectures, we considered varying heritability *h*^2^ and sparsity *s*, defined as the amount of variance explained by the genetic liability and the proportion of causal variants, respectively. For each setting of *h*^2^ and *s*, we simulated 100 different datasets. PRS models were derived by snpboost and evaluated by using various metrics regarding the predictive performance and the accuracy of the estimated coefficients. In detail, the predictive performance was measured by the MSEP and the *R*^2^ value defined as the squared correlation between the predicted and the true phenotype on the independent test set. To assess the computational efficiency we measured the computation time of the algorithm. The accuracy of the resulting estimates was evaluated by the number of included variants in the final model and the mean squared error (MSE) of the estimates as well as the true positive (TP) and true negative (TN) rate of the coefficients.

### Comparison to original *L*_2_-boosting in smaller settings

To analyse the performance of snpboost compared to the original component-wise *L*_2_-boosting algorithm [27], we used a single large batch with batch size *p*_batch_ = *p* in the snpboost algorithm on simulated data with reduced dimensionality. We then compared the results to the ones derived by using smaller batches in terms of predictive performance, computation time, mean squared errors of the estimated coefficients as well as true positive and true negative rates. The simulations were conducted for *n* = 20, 000 observations (10, 000 training set, 4, 000 validation set, 6, 000 test set) and *p* = 20, 000 variants as well as for varying degrees of heritability and sparsity. To obtain comparable results we chose a fixed number of boosting iterations independent of the batch size *p*_batch_ and a fixed learning rate *ν* = 0.1. For each simulation, 10 CPUs with 1 GB memory each were used.

Fig 2 displays the boxplots of each metric obtained after 1, 500 boosting iterations for heritability *h*^2^ = 50% and sparsity *s* = 0.1%, i.e. 20 influential variants. Incorporating batches did not largely affect the predictive performance in terms of *R*^2^ and MSEP nor the MSE of the coefficient estimates (MSE results not shown). However, different batch sizes do not always yield the same models as *L*_2_-boosting as can be seen by the number of variants included in the final model. The models resulting from a batch size of *p*_batch_ = 1000 tend to contain less variants than the one from original *L*_2_-boosting. This could be explained by the reduced search space in each boosting step. As a consequence, variants inside the batch that are already in the model are more often updated instead of including new variants outside of the batch. The fact that the other metrics remain almost constant suggests that either only variants with very small effects are not included when using a larger batch size or the variants that are updated are highly correlated with the ones not included. Furthermore, incorporating batches in the algorithm has a major effect on the computation time. To interpret the results shown in Fig 2 it is important to understand the two drivers of the computation time. On the one hand, it increases with the number of correlations that have to be calculated in each boosting step which explains the increased computation time of the original *L*_2_-boosting (i.e. a batch size of *p*_batch_ = 20, 000 and 20, 000 computed correlations in each boosting step) compared to smaller batch sizes such as *p*_batch_ = 100 and *p*_batch_ = 1000. On the other hand, reading the genotype data from disk when building the batches also increases the computation time leading to a higher computation time for smaller batches with *p*_batch_ = 10 for which more reads-from-disk have to be carried out. The varying computation times therefore reflect a trade-off between the number of correlations computed in each boosting step and the number of created batches.

**Fig 2.**
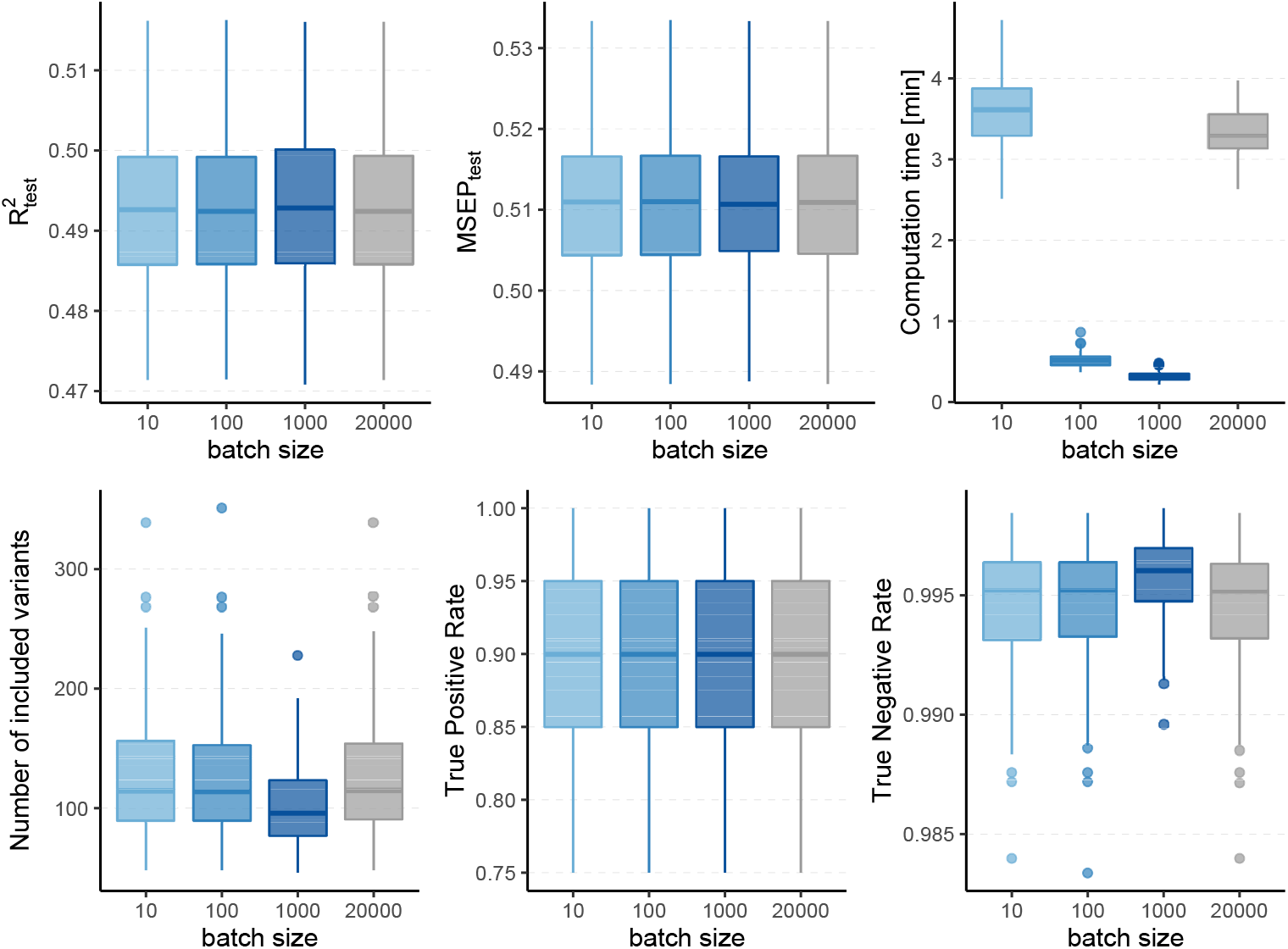
Comparison to original *L*_2_-boosting. Results of 100 simulated phenotypes with heritability *h*^2^ = 50% and sparsity *s* = 0.1% for *p* = 20, 000 variants and *n* = 20, 000 individuals (divided into 50% training, 20 % validation and 30% test set). Boxplots of the evaluation metrics obtained after 1, 500 boosting iterations are shown depending on the batch size. Batch size *p*_batch_ = 20, 000 corresponds to the original *L*_2_-boosting (shown in grey).

In summary, the incorporation of batches in the boosting algorithm did not affect the predictive performance of the model in our scenarios, while computation time was substantially reduced. However, snpboost does not always lead to the same models as the original *L*_2_-boosting algorithm, in particular in terms of the included variants and sparsity. The results for further settings with different heritability and sparsity were comparable and can be found in the supplemental material.

### Choice of hyperparameters for large-scale applications

The proposed snpboost algorithm includes various hyperparameters, namely the batch size *p*_batch_, the learning rate *ν*, the maximum number of boosting iterations per batch *m*_batch_, the maximum number of processed batches *b*_max_ and the stopping lag for the outer early stopping criterion *b*_stop_. In this section we discuss default values for the hyperparameters to facilitate the applicability of the algorithm in practice. The majority of these parameters do not need to be tuned but can be specified with reasonable default values, e.g. based on results from the literature and experience with the original boosting algorithm. For the remaining ones (*p*_batch_ and *b*_stop_) we examine how they influence the computational and predictive performance of snpboost in a simulation study.

The choice of the learning rate *ν* can be leaned on widely-used boosting algorithms. A rather small learning rate prevents boosting algorithms from overfitting on single base-learners and is therefore favorable regarding predictive performance. Nevertheless, a smaller learning rate will increase the number of needed boosting iterations to fit the full effect of the base-learners and simultaneously increase the algorithm’s computation time. Widely used R packages such as `mboost` [16, 32] and `xgboost` [33] use default learning rates of 0.1 and 0.3, respectively. As the effect of the learning rate will be comparable in the proposed adapted boosting algorithm, we decided to specify a fixed default value of *ν* = 0.1 in all our simulations. For the batch-related hyperparameters we varied the batch size *p*_batch_ over a range of possible values namely *p*_batch_ ∈ {10, 100, 1000, 5000} to analyse its effect. For each batch we allow a maximum number of boosting iterations *m*_batch_ equivalent to the batch size *p*_batch_. Since we specified the learning rate with a rather small fixed value and due to the correlation-based early stopping criterion, this choice should prevent the algorithm from overfitting on one batch. If one or more variants inside the batch are still among the most influential ones out of all variants they will also be included in the next batch. For the outer stopping criterion we specified a large maximum number of batches *b*_max_ = 20, 000 to ensure that the algorithm terminates even in case the MSEP on the validation set has not decreased for *b*_stop_ batches. Since we do not want the algorithm to stop too early and simultaneously minimize the computation time, in our simulations we consider the choices *b*_stop_ = 2 and *b*_stop_ = 10.

We then fitted PRS models using snpboost with the previously described hyperparameters. For the computations we used 10 CPUs with 2 GB RAM each.

The results for simulated phenotypes with 10% and 50% heritability are shown in Fig 3 and Fig 4. Results for further degrees of heritability can be found in the supplement. Independently of the heritability and the sparsity of the simulated data, the predictive performance was not affected in our settings by varying batch sizes in terms of *R*^2^ and MSEP. However, the computation time differed crucially, resulting in considerably higher values for rather small (*p*_batch_ = 10) or rather large (*p*_batch_ = 5000) batches. Furthermore, larger batches led to a higher number of included variants in the final model. This effect was stronger for phenotypes which have a less sparse genetic architecture and associated with a later stopping of the algorithm, i.e. more boosting steps were required to derive the final model. A higher number of variants in the final model was associated with a slightly higher MSE of the coefficients as well as higher true positive rates on the one hand but also smaller true negative rates on the other hand. As expected, a higher *m*_stop_ increased the computation time of the fitting process for all batch sizes. In contrast, there was no considerable effect on the predictive performance. However, *b*_stop_ = 2 and *b*_stop_ = 10 had an impact on the coefficient estimates as can be seen in Fig 4, e.g. by a tendency to include more variants in the model when choosing *b*_stop_ = 10. This tendency was only apparent for batch sizes *p*_batch_ < 1000, suggesting that for larger batches the choice of *b*_stop_ is only of minor importance for both, prediction performance and coefficient estimates. The results clearly indicate that a more favorable signal-to-noise ratio (i.e. a higher heritability) and less influential variants (i.e. a higher sparsity) are in general beneficial for the performance of our approach. For phenotypes with a sparser genetic architecture, the considered evaluation metrics tended to show less variability.

**Fig 3.**
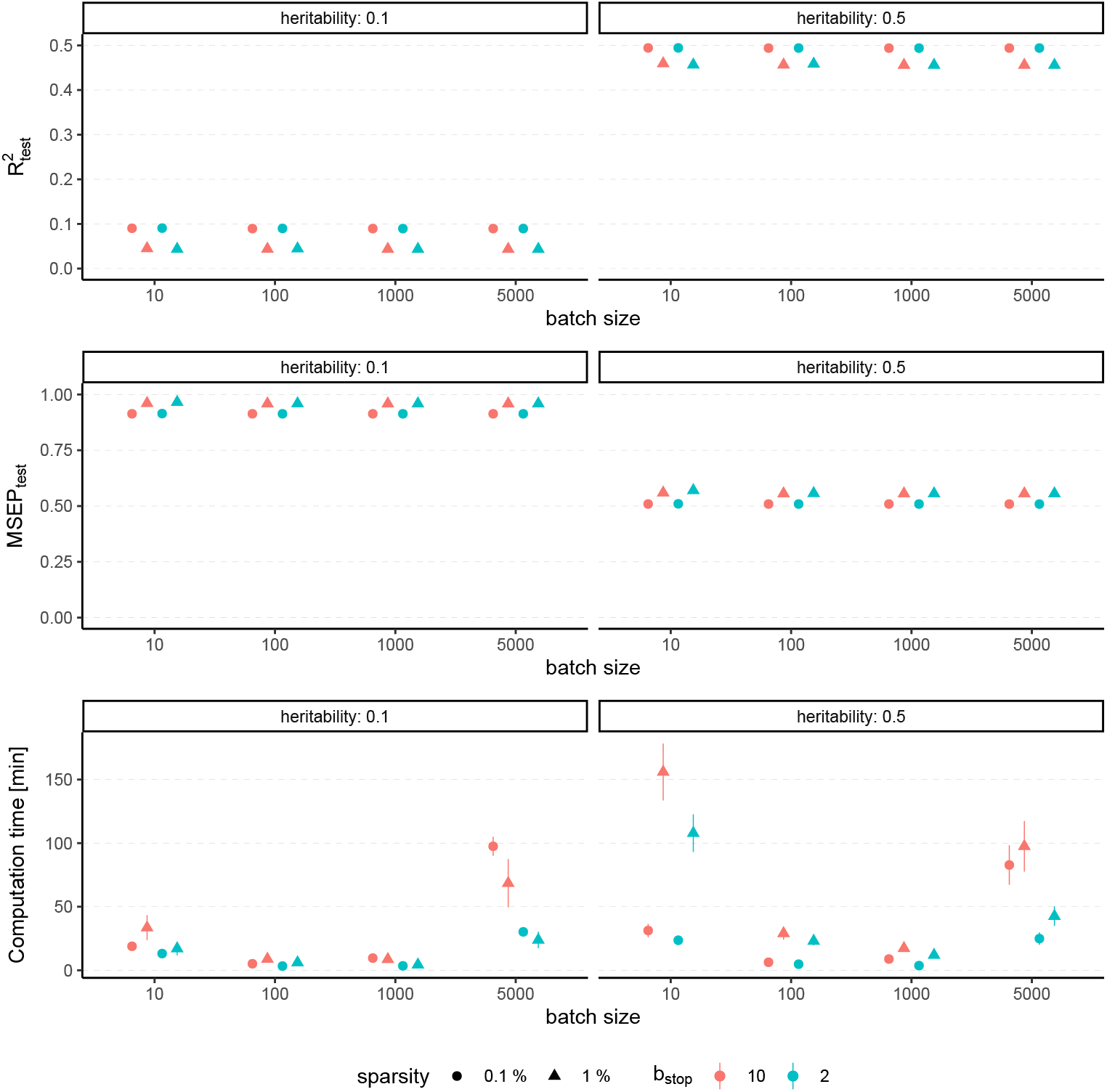
Predictive performance for varying batch size and stopping criteria. Results of 100 simulated phenotypes with heritability *h*^2^ ∈ {10%, 50%}, sparsity *s* ∈ {0.1%, 1%}and *b*_stop_ ∈ {2, 10}for *p* = 100, 000 variants and *n* = 100, 000 individuals (divided into 50% training, 20 % validation and 30% test set). Mean and standard deviation of the evaluation metrics are shown depending on the batch size.

**Fig 4.**
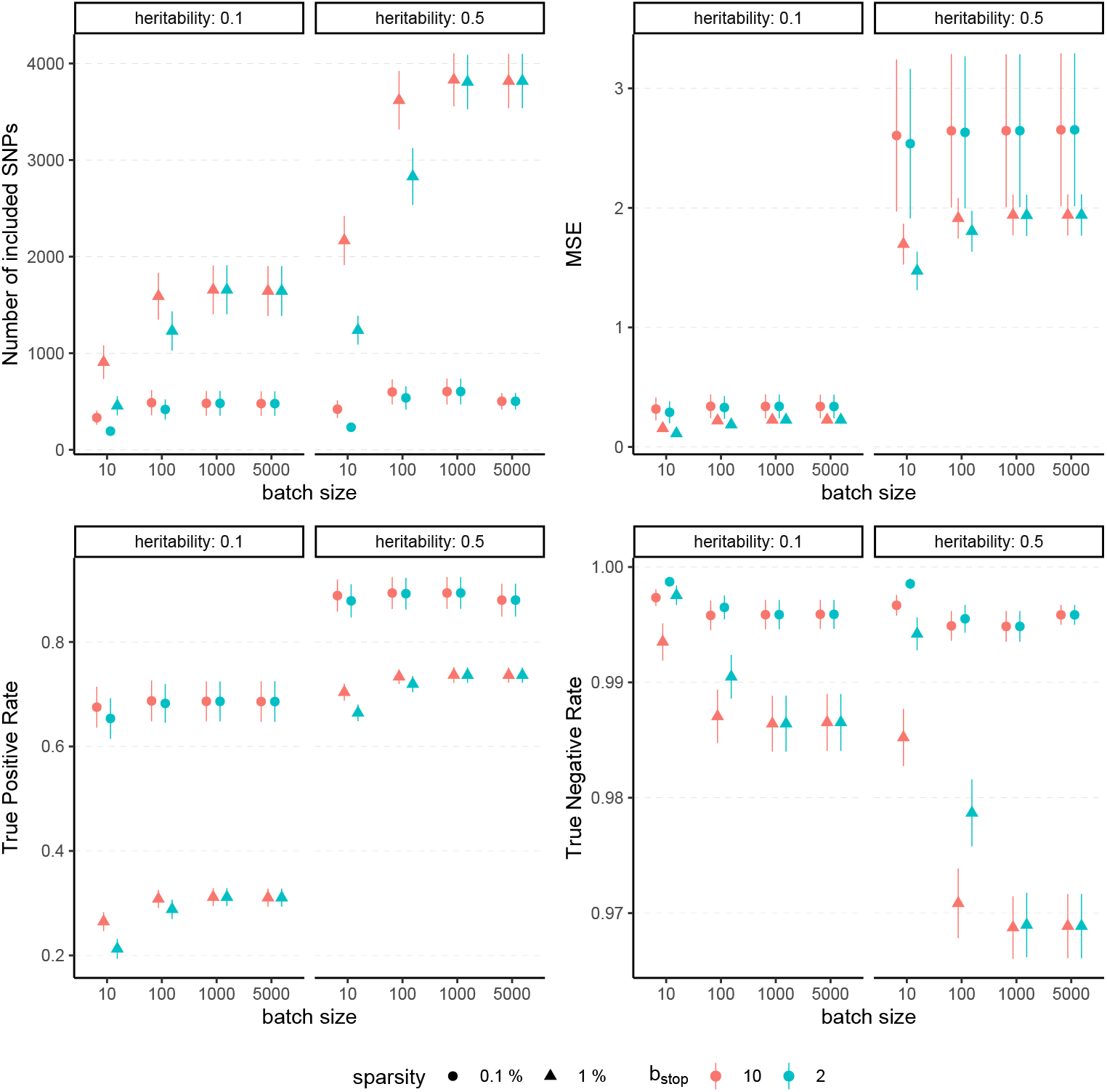
Evaluation metrics of the estimated coefficients for varying batch size and stopping criteria. Results of 100 simulated phenotypes with heritability *h*^2^ ∈ {10%, 50%}, sparsity *s* ∈ {0.1%, 1%}and *b*_stop_ ∈ {2, 10} for *p* = 100, 000 variants and *n* = 100, 000 individuals (divided into 50% training, 20 % validation and 30% test set). Mean and standard deviation of the evaluation metrics are shown depending on the batch size.

In summary, the choice of the hyperparameters had no major influence on the predictive performance measures *R*^2^ and MSEP but on the computation time, which was lowest for medium size batches (100 ≤ *p*_batch_ ≤ 1000). The accuracy of the coefficient estimates measured via MSE, TP and TN rate varied with the batch size, as larger batches tended to lead to more (true positive) variants included in the final model, but also to a slightly higher MSE and a smaller TN rate. While the differences in MSE, TP and TN rate were only small, smaller batches yielded sparser models in particular for phenotypes with a high heritability.

To conclude, batch sizes of 100 ≤ *p*_batch_ ≤ 1000 seem to be the most favorable regarding the computation time and the other evaluation metrics. We propose a batch size of *p*_batch_ = 1000 as the default value because the results suggest less dependency on the *b*_stop_ parameter than for a batch size of 100 variants. Accordingly, we recommend a default value of *b*_stop_ = 2 to keep the computation time as low as possible. In practice, genotype data often contain more than 100, 000 variants, which further supports the choice of *p*_batch_ = 1000 with regard to the computation time. Although our simulation study suggests that those default values should provide reasonable results in most cases, it is recommendable to take the genetic architecture of the examined phenotype as well as the main aim of the analysis into account. Phenotypes with a high expected heritability might be better fitted by using smaller batches, while for phenotypes with many causal variants larger batches might be favorable to increase the TP rate. If one is interested in extremely sparse models identifying only the most-informative variants one could also try to use smaller batches to avoid an overestimation of the number of causal variants.

## Application to the UK Biobank

We applied our proposed method on data from the UK Biobank resource under Application Number 81202. Besides the validation of the results from the previous section, we compared our boosting models fitted via the proposed snpboost approach to the ones derived by fitting the lasso via the BASIL algorithm implemented in the `snpnet` package [11], which have been shown to outperform commonly-used PRS models based on univariate summary statistics for various traits.

The UK Biobank (UKBB, [30]) is a large-scale prospective cohort study including more than half a million participants from the United Kingdom aged between 40 and 69 years. The database comprises genome-wide genotype data of each individual as well as various in-depth phenotypic information such as biological measurements as well as blood and urine biomarkers. The data have been collected since 2006 and are continually updated with follow-up data.

Our aim is to estimate PRS for various phenotypes, covering several heritability and sparsity levels. The heritability of a trait is an upper bound for the predictive performance based on genotype information. Thus, we used the analyses of Tanigawa et al. [34] as a proxy and specifically considered five appropriate continuous phenotypes: standing height in cm (UKBB field 50), LDL-cholesterol in mmol/l (UKBB field 30780), blood glucose level in mmol/L (UKBB field 30740), lipoprotein A in nmol/L (UKBB field 30790) and BMI in kg/m^2^ (UKBB field 21001).

Height and BMI are quantitative traits with a relatively high heritability and a rather polygenic structure. Twin-studies estimated a heritability of approximately 69% for height and 42% for BMI after adjusting for covariates [35]. For a long time, genetic models could not explain this estimated heritability, a phenomenon known as “missing heritability” [36, 37]. More recent studies have indicated that this may be primarily due to many influential common variants with small effect sizes [38–40] underlining the high polygenicity of those traits. In contrast to this, the distribution of the biomarker lipoprotein A, which is a strong risk factor for coronary heart disease, is mainly explained by variants in the LPA gene on chromosome 6 [41]. Thus, we expect a sparse PRS with a relatively high prediction accuracy for this trait. For LDL-cholesterol it is known that it is associated with several genes such as LDLR and PCSK9 [42, 43]. Therefore, we expect signal in several genomic regions. Recent studies compared different approaches including the lasso to derive PRS, and found that multivariable methods can reach a predictive performance of up to 20% [12, 34]. As in previous works [44], we adjusted the measured LDL-cholesterol value by a factor of 0.684 for individuals taking statins lowering the blood lipid. For blood glucose we are not aware of a genetic impact and also Tanigawa et al. [34] found the genetic background only explaining a small fraction (less than 2%) of the biomarker’s variance.

Out of the over 500,000 individuals from UK Biobank we filtered for individuals with self-reported white British ancestry (UKBB field 21000) and available data for all chosen phenotypes, resulting in *n* = 284, 342 observations. Additionally, the covariates age and sex as well as the first ten principal components of the genotype matrix are available. We randomly divided the data set into training (*n_train_* = 170, 557), validation (*n_val_* = 56, 801) and test set (*n_test_* = 56, 984). We used genome-wide genotype data and filtered for variants with a genotyped rate of at least 90% and a minor allele frequency of at least 0.1%, resulting in *p* = 562, 684 genetic variants. Missing genotypes are imputed by the corresponding mean of the complete observations.

For both the boosting and lasso approaches, we first estimated a PRS using only the genotyped variants as predictors. We used the training set to fit the model and the validation set to simultaneously monitor the predictive performance for choosing the main tuning parameters of the algorithms (i.e., the number of iterations for boosting and the penalty parameter for the lasso). To fit the lasso we used the R package `snpnet` [11] with the provided default hyperparameters. Following the results of our simulation study, for the snpboost algorithm we chose a batch size of *p*_batch_ = 1, 000 variants, a learning rate of *ν* = 0.1 and an outer stopping lag of *b*_stop_ = 2 batches. Using the resulting estimated 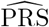 we fitted two linear models, namely the first one (*M*_PRS_) incorporating only the PRS as a single predictor variable:

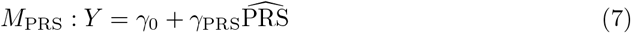

and the second one (*M_f_*) including the first ten principle components, sex and age as additional covariates:

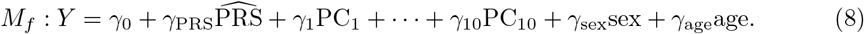

To measure the actual benefit in accuracy of including a PRS in the prediction model, we also fitted a model including only covariates (*M_c_*):

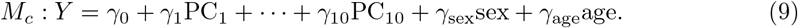

Finally, we also included the covariates in the fitting process to derive the PRS, corresponding to the model *M_PRS,c_*:

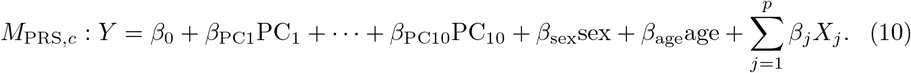

All models were evaluated on the independent test set and compared with respect to their predictive performance, computational efficiency and sparsity. To measure the predictive performance we used the *R*^2^ value on the test set given by the squared correlation between the observed and predicted phenotypes as well as the root mean squared error of prediction (RMSEP). The computational efficiency was measured as the computation time in minutes of the respective algorithm and the sparsity is given by the number of included variants in the final PRS (*N* variants). All computations were conducted on a computer cluster with 16 CPUs and 2 GB RAM each.

The results for all phenotypes are given in Table 2 and Table 3. Overall, snpnet and snpboost yield comparable results regarding the predictive performance, without one approach being consistently superior to the other. Both the resulting *R*^2^ and RMSEP are very close. Furthermore, the shown *R*^2^ values are in line with previously reported *R*^2^ resulting from snpnet, which has been shown to be highly competitive to various other (univariate) PRS methods [11, 34]. The PRS estimated via snpnet and snpboost both clearly increase the predictive performance compared to the covariates-only model *M_c_* for all shown phenotypes. With respect to sparsity, our boosting approach tends to select less variants (on average 16% less variants compared to the lasso). The computation time of both approaches is highly dependent on the genetic architecture, i.e. the heritability and sparsity of the phenotype. In particular, a higher and more polygenic signal tends to lead to longer computation times. In case of fitting the PRS based solely on the genotype data and including the covariates in a subsequent linear model, snpboost tends to be faster than snpnet; however, the computation times for snpboost increase substantially when including covariates in the fitting process for LDL-cholesterol and height. This is partly due to more coefficients being fitted and updated in each boosting step and partly due to larger PRS models resulting from more boosting steps. Nevertheless, the models are still fitted in reasonable time using our batch-based approach. As described in Hepp et al. [26], boosting is generally expected to be slower than the lasso, which can only be observed for less sparse models in the examined phenotypes. In general, the model *M*_PRS,*c*_ outperforms the model *M_f_* regarding the predictive performance, implying that including the covariates already in the fitting of the PRS is favorable regarding the detection of the genetic signal. However, the effect is only substantial for phenotypes with a high association to covariates (i.e. height). Furthermore, the model *M*_PRS,*c*_ tends to select more variables than estimating the PRS based only on the genotypes (*M*_PRS_) and the computation time is considerably increased when using the snpboost approach. Therefore, it might be advisable to only consider the covariates in the fitting process if there is a large association already in the covariates-only model.

**Table 2.**
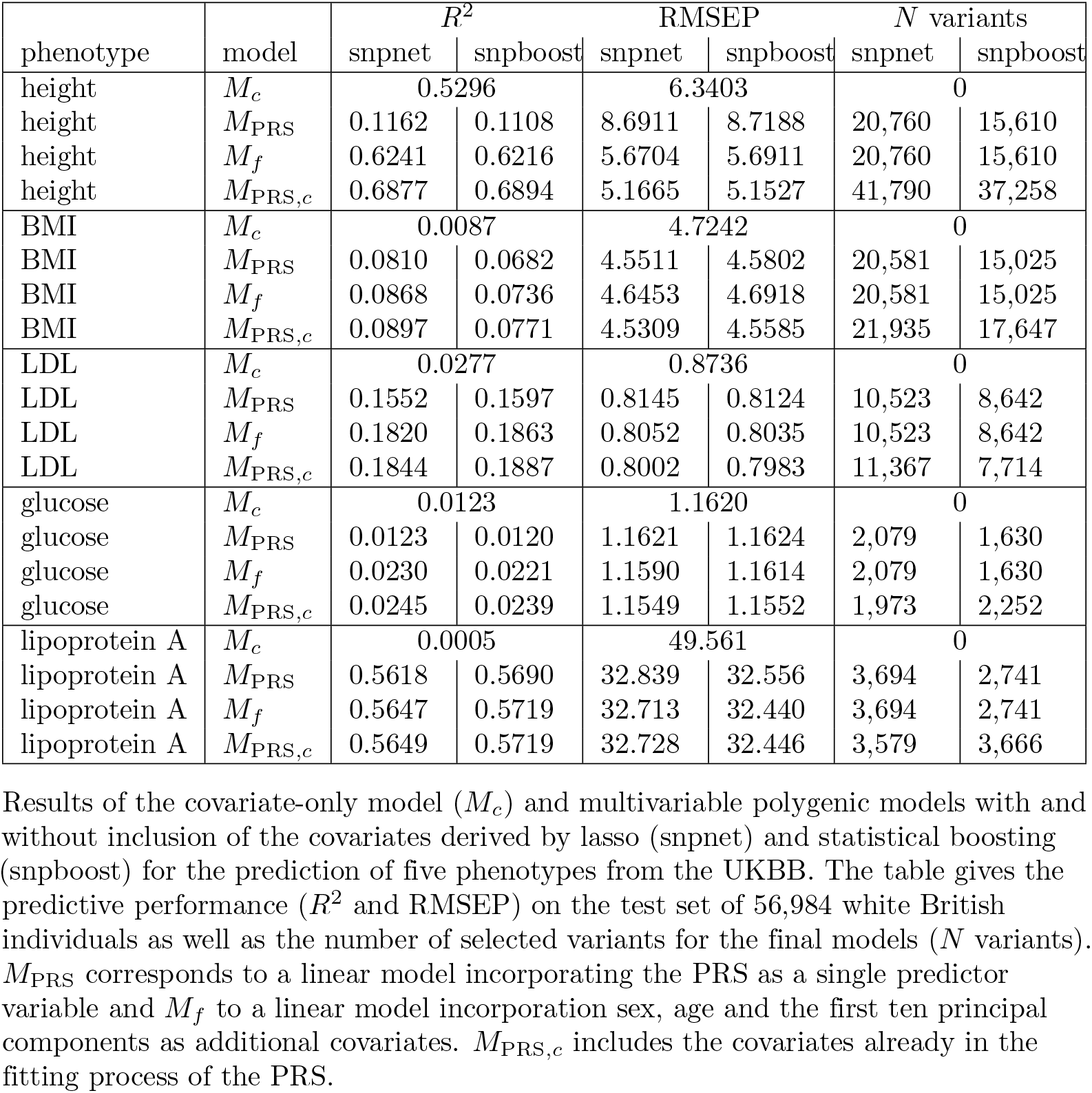
Comparison of predictive performance of snpnet and snpboost for five phenotypes from the UKBB.

**Table 3.** Comparison of computational efficiency of snpnet and snpboost on five phenotypes from the UKBB.

| phenotype | model | computation time in minutes |  |
| --- | --- | --- | --- |
|  |  | snpnet | snpboost |
| height | $M_{\text{PRS}}$ | 121.11 | 90.73 |
| height | $M_{\text{PRS},c}$ | 121.92 | 377.61 |
| BMI | $M_{\text{PRS}}$ | 70.29 | 76.47 |
| BMI | $M_{\text{PRS},c}$ | 84.90 | 220.37 |
| LDL | $M_{\text{PRS}}$ | 65.15 | 45.47 |
| LDL | $M_{\text{PRS},c}$ | 52.79 | 63.33 |
| glucose | $M_{\text{PRS}}$ | 24.47 | 11.89 |
| glucose | $M_{\text{PRS},c}$ | 23.51 | 19.23 |
| lipoprotein A | $M_{\text{PRS}}$ | 40.43 | 20.35 |
| lipoprotein A | $M_{\text{PRS},c}$ | 42.68 | 39.15 |

Computational times of the algorithms snpnet and snpboost for multivariable polygenic models with and without inclusion of the covariates for the prediction of five phenotypes from the UKBB. $M_{\text{PRS}}$ corresponds to the application of the algorithms without including covariates and $M_{\text{PRS},c}$ to the inclusion of the covariates sex, age and the first ten principal components. The experiments were run on 16 CPUs with 2GB RAM each.

Fig 5 displays the absolute values of the resulting estimated non-zero coefficients for LDL-cholesterol for the boosting and lasso approaches. Both tend to detect variants with higher effect sizes in the same genetic regions, e.g. at chromosome 2 and chromosome 19. In total, there are 3, 675 genetic variants that are present in both PRS, out of 8, 642 variants selected by snpboost and 10, 523 variants selected by snpnet. While snpboost results in less variants, the included variants have larger effect sizes and less variants with very small effect sizes close to zero are included in the model. Supplementary Fig S5 displays the coefficients again with shared variants marked in black. All SNPs with comparably high effect sizes in the snpnet PRS are included in both models but the snpboost PRS incorporates further SNPs with stronger effects. The results are similar for the other phenotypes and included in the supplementary material. In conclusion, the snpboost PRS tends to include less variants in total, but more variants with comparably high effect sizes corresponding to less shrinkage for the variants included in the model compared to the lasso.

**Fig 5.**
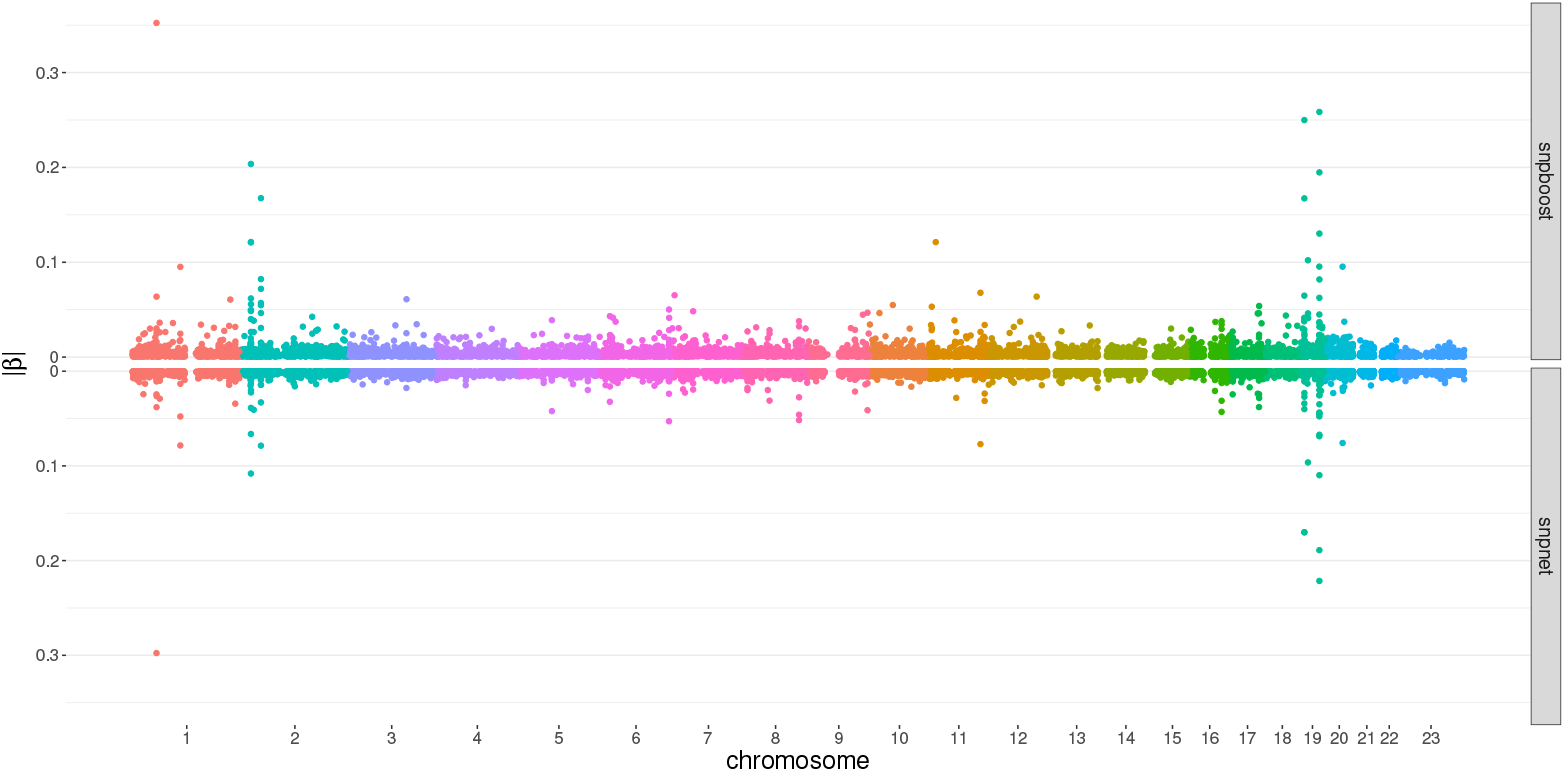
Absolute values of coefficient estimates for PRS models for LDL-cholesterol derived by boosting (snpboost) and lasso (snpnet) in dependence of the genomic position of the variants.

## Discussion

In this work we have proposed a new methodological framework to derive multivariable PRS models via applying a statistical boosting approach directly on genotype data. Currently, PRS are most often built based on summary statistics from GWAS that were estimated by simple and univariate linear regression models [5]. This methodologically simple approach is mainly justified by the computational hurdle resulting from the ultra-highdimensionality of the the genotype data. However, recently published works provided methods to enable statistical modelling by penalized multivariable regression approaches on genotype data [10, 11]. Qian et al. demonstrated that lasso-based PRS were able to outperform several PRS derived by methods based on summary statistics [11]. First approaches to apply statistical boosting on genotype data employed a three-step-approach to fit multivariable PRS [12]: first, variants are pre-filtered based on their univariate associations with the examined phenotype. Second, statistical modelling and variable selection approaches such as AdaSub [20] and boosting with probing [18] are used to identify the informative variants on blocks of variants in LD. Finally, a multivariable PRS based on the selected variants is constructed via component-wise *L*_2_-boosting [16]. While this approach yielded particularly sparse models and could compete with common methods like clumping and thresholding [6], lasso via `snpnet` clearly yielded more accurate results regarding the predictive performance which is usually the main objective of PRS modelling.

In the present article we introduced the boosting algorithm snpboost that works on smaller batches of variants similar to the BASIL algorithm. Our framework enables the application of statistical boosting directly on the complete original genotype data. In a smaller but still high-dimensional simulation setting, we were able to show that the adapted boosting algorithm yields similar performance to the original component-wise *L*_2_-boosting, indicating that we do not lose predictive performance due to the incorporation of batches. In a further setting with more realistic dimensionality we have derived appropriate default values for the application of snpboost on large-scale data. We have shown that the specified default values resulted in reasonable models in most cases and also have given advice to adapt them based on the genetic architecture of the examined phenotype and the specific research questions.

We applied the newly proposed snpboost algorithm on large-scale genotype data of the UKBB. In particular, we have compared the performance of snpboost to the one achieved by the lasso via snpnet, which has been shown to outperform many classical PRS [11]. Our results indicate that the snpboost algorithm leads to PRS models that are highly competitive to lasso-based PRS models in both predictive performance and computation time. Although it might have been expected that the computation time would be higher for statistical boosting than for lasso [26], our approach had a tendency to result in sparser models. These sparser models correspond to an earlier stopping of the algorithm which reduces the computation time of boosting. The incorporation of further covariates such as age, sex and principal components in the fitting process of the PRS resulted in increased computation times for some phenotypes, particularly for height. However, in such cases, the boosting algorithm yielded an improved predictive performance with larger numbers of included variants. This illustrates that sparsity is not always favorable in regards of predictive performance.

Our analyses show that there is a large overlap of the chosen variants by lasso and boosting, in particular regrading the variants with high estimated effect sizes. However, boosting has been found to include less variants in the final model and to induce less shrinkage on the effect estimates compared to the lasso. In clinical practice, a sparser PRS model might be of particular interest if the aim is not only prediction but also the identification of risk loci in the genome. In fact, functional annotations of the selected variants can better elucidate the underlying biological mechanisms influencing the analyzed trait. Thus, statistical boosting might be one way to yield more biologically interpretable PRS models.

Despite the presented promising results, we inherited also some limitations from statistical boosting. In contrast to classical regression methods, boosting does not provide closed formulas for standard errors of effect estimates or confidence intervals that could be used for inference. Furthermore, as mentioned before, statistical boosting is in general associated with a slightly higher computational complexity compared to methods such as the lasso [26] and has a known tendency to include too many variables in low-dimensional settings [19, 45]. Our results suggest that the incorporation of batches substantially reduced the computational time. Additionally, the reduction of the search space in each boosting step might partially prevent the algorithm from selecting too many variables. However, the implementation of the batch-based statistical boosting in snpboost is currently limited to continuous phenotypes and linear base-learners, each corresponding to one genetic variant.

In future research we want to exploit the modular structure of boosting which makes the algorithm very flexible to extend. We will incorporate different loss functions to extend the snpboost framework to be applicable also to binary and time-to-event data. To account for the uncertainty in the prediction, one could also construct subject-specific prediction intervals based on quantile regression [46]. Besides extending the approach via new loss functions, one could also adapt the base-learners in various ways. For example, base-learners could be adapted to take into account different models of inheritance beside the classical additive component typically used in the polygenic models, such as dominant and recessive hereditary schemes. Further possibilities include the extension of the set of possible base-learners to interactions between variants and variant-covariate interactions meaning that the boosting algorithm can be adapted to model gene-environment interactions as well as epistatic effects across variants which can play a relevant role in biological phenotypes [47]. Besides those methodological extensions of our proposed snpboost framework, future research will also focus on the practical application of PRS derived by our framework. An important aspect of PRS research is the transferability of PRS models to different ethnicities, as PRS are often derived on cohorts of European ancestry and a substantial loss of predictive performance is observed when applied on further cohorts with different ethnicities [48, 49]. Previous studies have indicated that sparser models may contribute to overcome this issue [12] and it is of particular interest to examine the transferability of PRS derived by statistical boosting.

## Acknowledgments

The work on this article was supported by the Deutsche Forschungsgemeinschaft (DFG, grant number 428239776, MA7304/1-1).

This research has been conducted using the UK Biobank resource under application number 81202.

