## Supplemental Figures for "Boosting polygenic risk scores"

### Supporting information

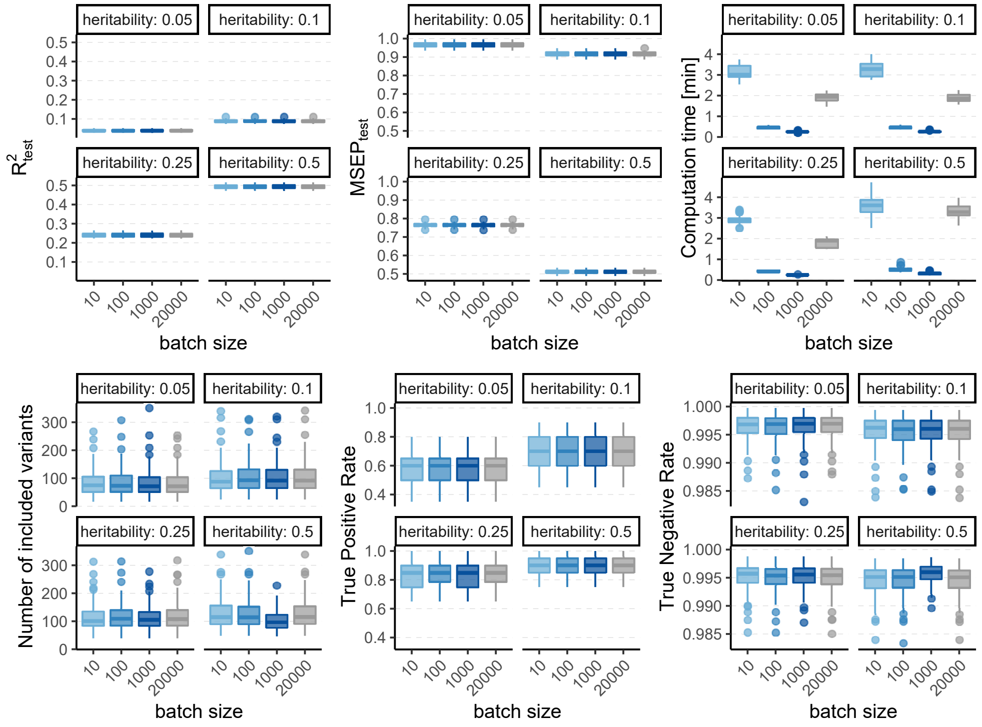


**S1 Fig. Results of 100 simulated phenotypes with varying heritability and sparsity $s = 0.1\%$ for $p = 20,000$ variants and $n = 20,000$ individuals (divided into 50% training, 20 % validation and 30% test set).** Boxplots of the evaluation metrics obtained after $1,500$ boosting iterations are shown depending on the batch size. Batch size $p_{\text{batch}} = 20,000$ corresponds to the original $L_2$-boosting (shown in grey).

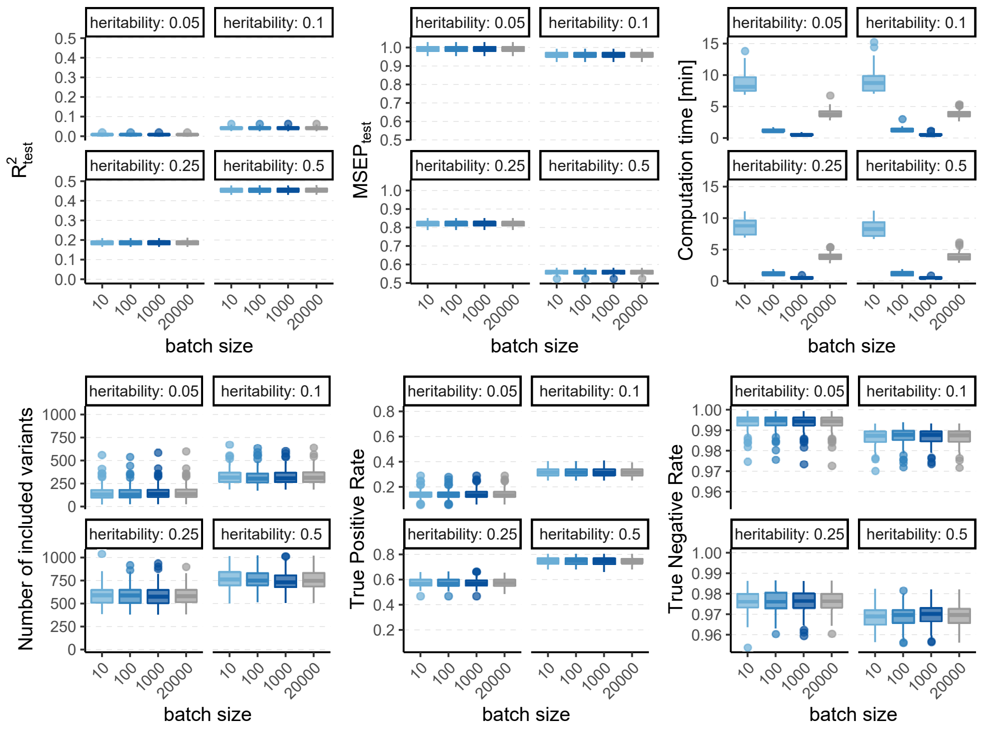


**S2 Fig. Results of 100 simulated phenotypes with varying heritability and sparsity $s = 1\%$ for $p = 20,000$ variants and $n = 20,000$ individuals (divided into 50% training, 20 % validation and 30% test set).** Boxplots of the evaluation metrics obtained after 3,500 boosting iterations are shown depending on the batch size. Batch size $p_{\text{batch}} = 20,000$ corresponds to the original $L_2$-boosting (shown in grey).

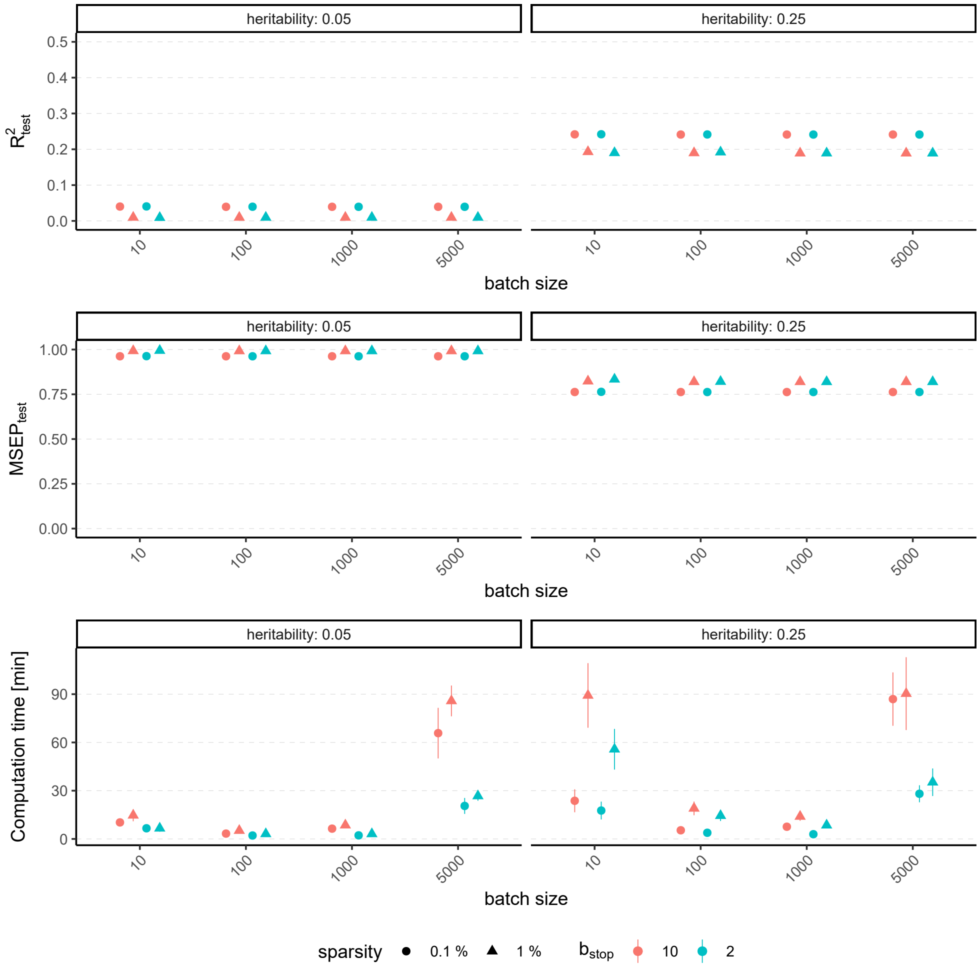


**S3 Fig. Results of 100 simulated phenotypes with heritability $h^2 \in \{5\%, 25\%\}$, sparsity $s \in \{0.1\%, 1\%\}$ and $b_{stop} \in \{2, 10\}$ for $p = 100,000$ variants and $n = 100,000$ individuals (divided into 50% training, 20 % validation and 30% test set).** Mean and standard deviation of the evaluation metrics are shown depending on the batch size.

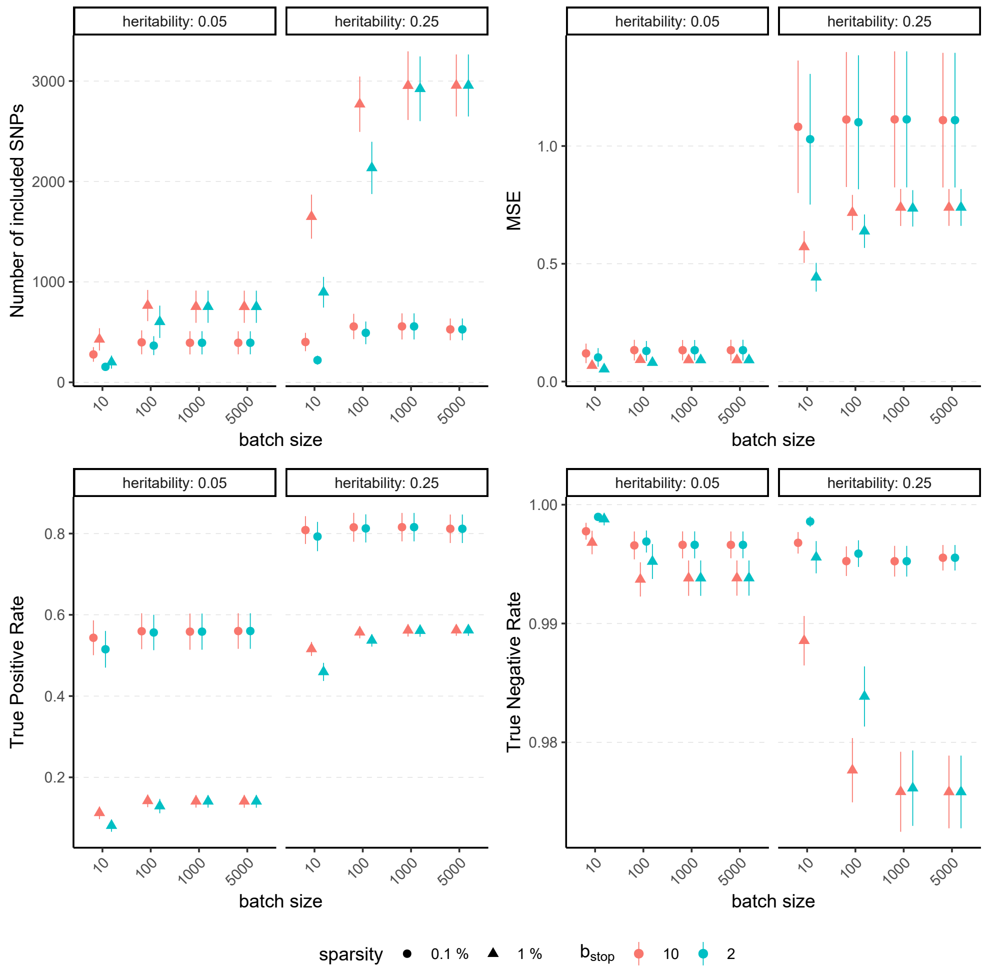


**S4 Fig. Evaluation metrics of the estimated coefficients for 100 simulated phenotypes with heritability $h^2 \in \{5\%, 25\%\}$, sparsity $s \in \{0.1\%, 1\%\}$ and $b_{\text{stop}} \in \{2, 10\}$ for $p = 100,000$ variants and $n = 100,000$ individuals (divided into 50% training, 20 % validation and 30% test set).** Mean and standard deviation of the evaluation metrics are shown depending on the batch size.

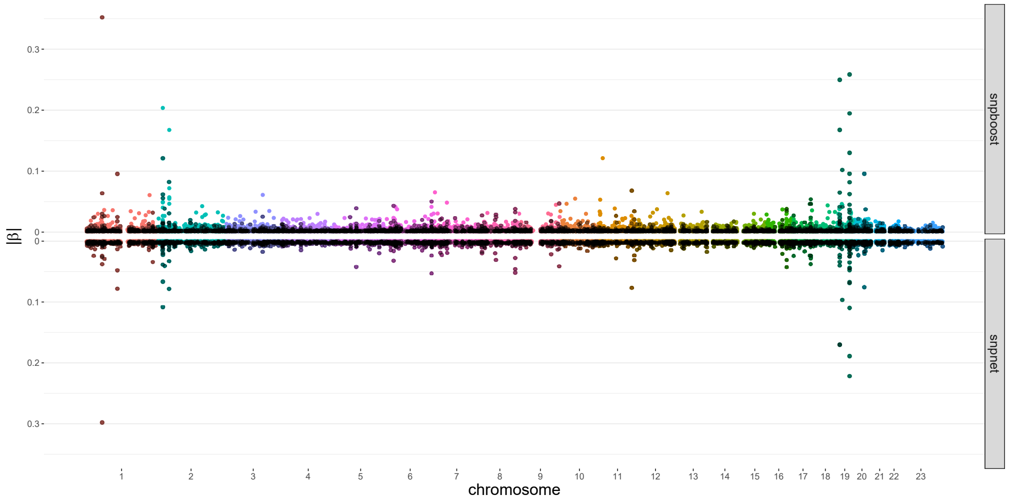


**S5 Fig. Absolute values of coefficient estimates for PRS models for LDL-cholesterol derived by boosting (snpboost) and lasso (snpnet) shown in dependence of the genomic position of the variants. Variants that are included in both models are marked in black.**

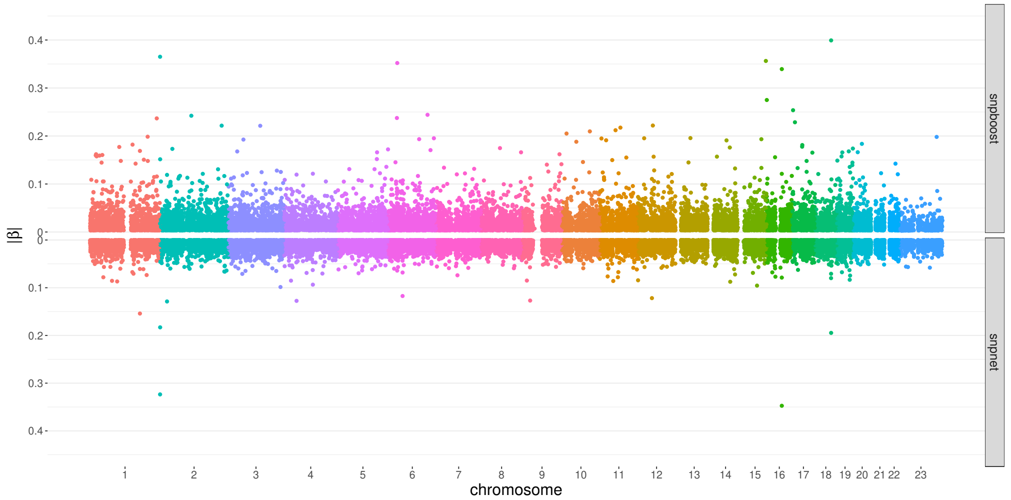


**S6 Fig. Absolute values of coefficient estimates for PRS models for BMI derived by boosting (snpboost) and lasso (snpnet) shown in dependence of the genomic position of the variants.**

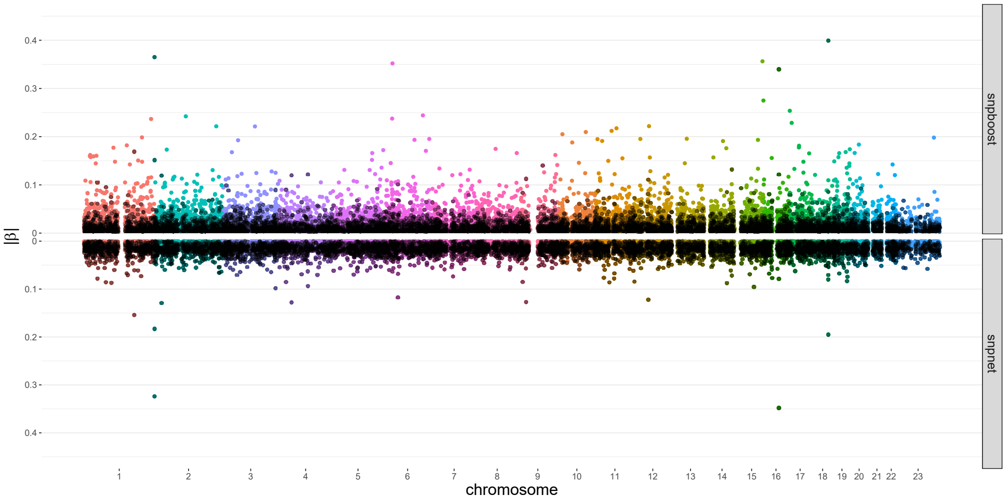


**S7 Fig. Absolute values of coefficient estimates for PRS models for BMI derived by boosting (snpboost) and lasso (snpnet) shown in dependence of the genomic position of the variants. Variants that are included in both models are marked in black.**

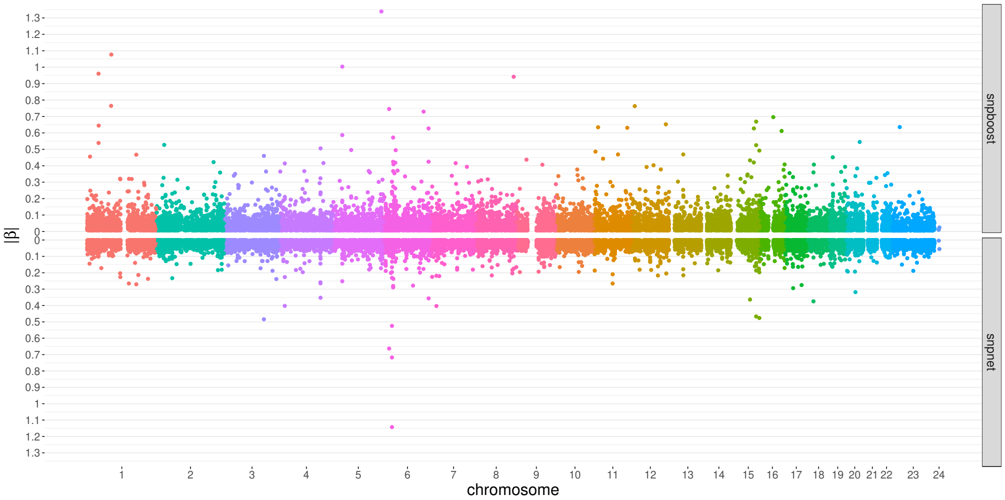


**S8 Fig. Absolute values of coefficient estimates for PRS models for height derived by boosting (snpboost) and lasso (snpnet) shown in dependence of the genomic position of the variants.**

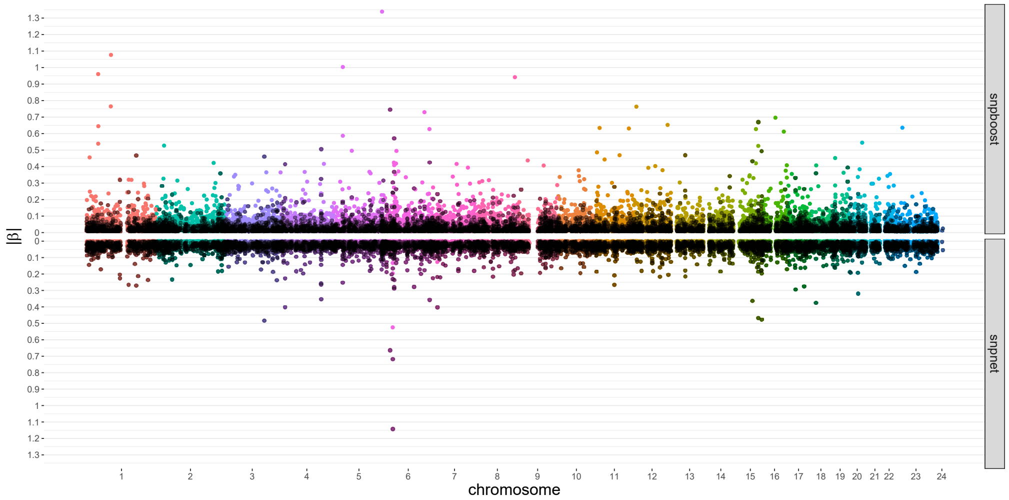


**S9 Fig. Absolute values of coefficient estimates for PRS models for height derived by boosting (snpboost) and lasso (snpnet) shown in dependence of the genomic position of the variants. Variants that are included in both models are marked in black.**

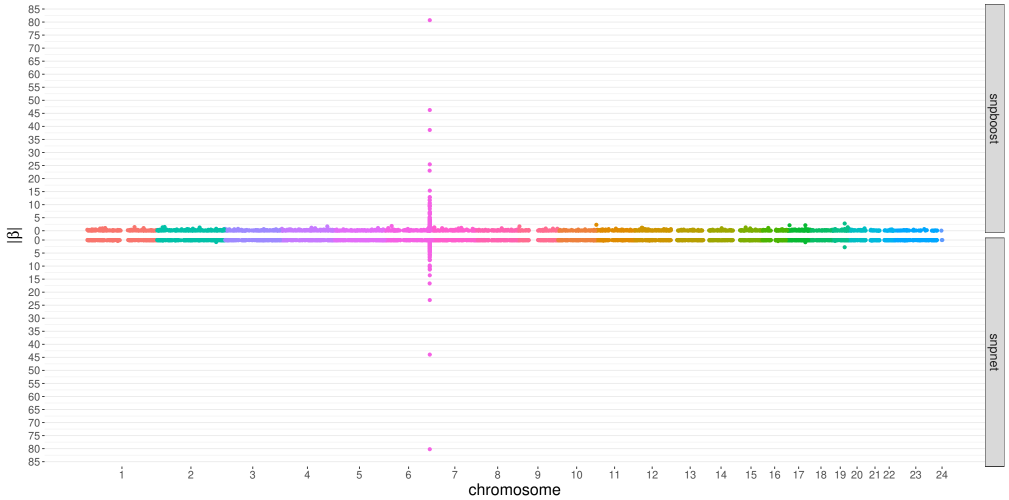


**S10 Fig. Absolute values of coefficient estimates for PRS models for lipoprotein A derived by boosting (snpboost) and lasso (snpnet) shown in dependence of the genomic position of the variants.**

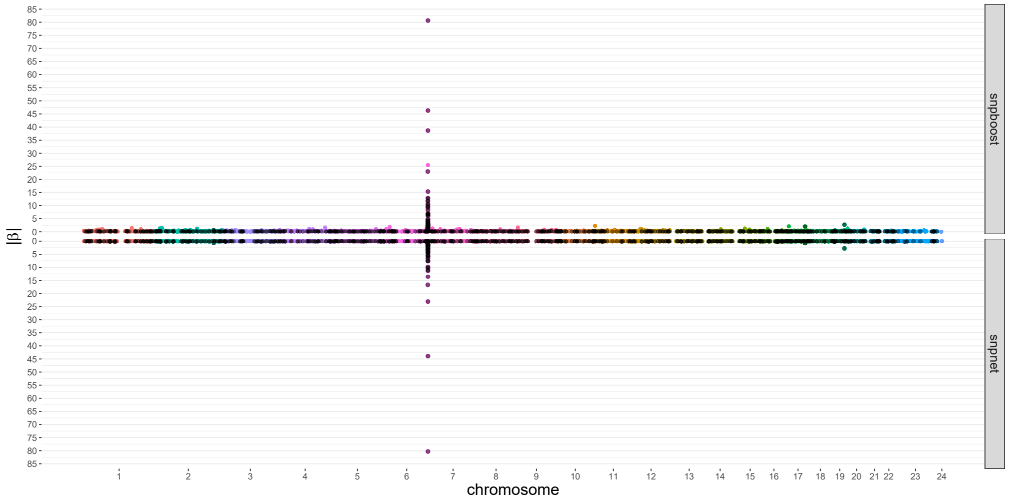


**S11 Fig. Absolute values of coefficient estimates for PRS models for lipoprotein A derived by boosting (snpboost) and lasso (snpnet) shown in dependence of the genomic position of the variants. Variants that are included in both models are marked in black.**

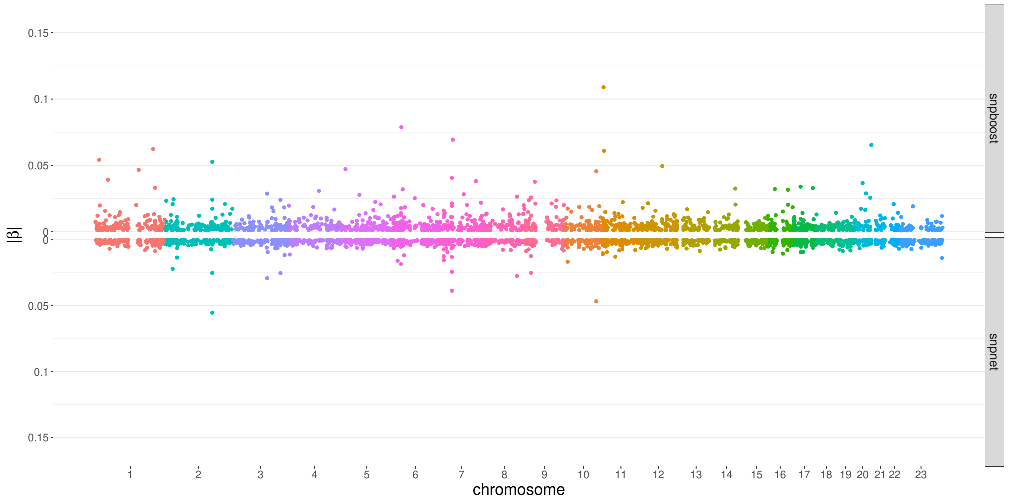


**S12 Fig. Absolute values of coefficient estimates for PRS models for glucose derived by boosting (snpboost) and lasso (snpnet) shown in dependence of the genomic position of the variants.**

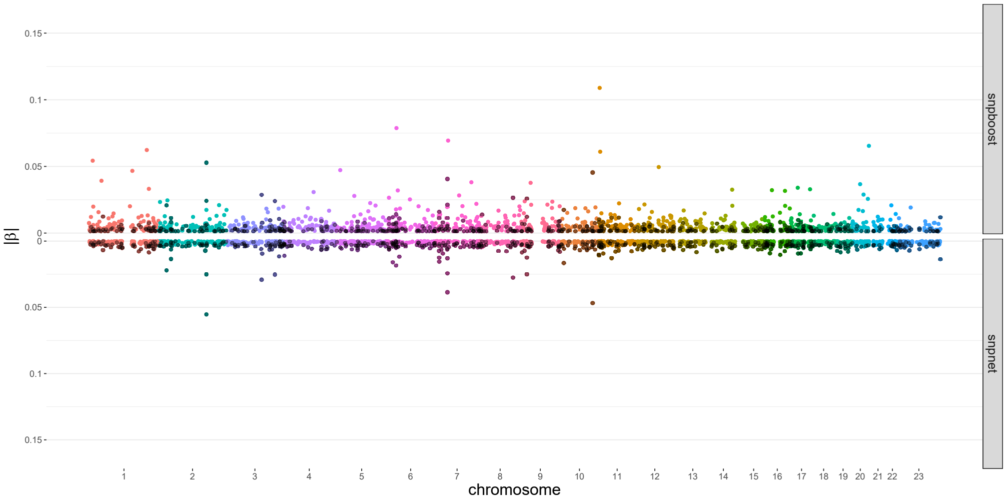


**S13 Fig. Absolute values of coefficient estimates for PRS models for glucose derived by boosting (snpboost) and lasso (snpnet) shown in dependence of the genomic position of the variants. Variants that are included in both models are marked in black.**
